# Refining sequence-to-activity models by increasing model resolution

**DOI:** 10.1101/2025.01.24.634804

**Authors:** Nuria Alina Chandra, Yan Hu, Jason D. Buenrostro, Sara Mostafavi, Alexander Sasse

**Affiliations:** Paul G. Allen School of Computer Science and Engineering, University of Washington, Seattle 98195, WA, USA; Gene Regulation Observatory, Broad Institute of MIT and Harvard, Cambridge 02142, MA, USA; Department of Stem Cell and Regenerative Biology, Harvard University, Cambridge, 02138, MA, USA; Canadian Institute for Advanced Research, Toronto, MG51ZB, ON, Canada; Center for Synthetic Genomics, Center for Molecular Biology Heidelberg (ZMBH), Heidelberg University, Heidelberg, 69120, Germany

**Keywords:** ATAC-seq, chromatin, gene regulation, deep learning, sequence-to-function models

## Abstract

Decoding the cis-regulatory syntax that controls gene expression is essential for improving our understanding of cell differentiation and disease. To identify regulatory motifs and their regulatory syntax, deep learning based sequence-to-activity (S2A) models learn transcription factor binding motifs and their combinations from DNA sequence by modeling measured chromatin accessibility. Previously, we developed AI-TAC, a S2A model that predicts chromatin accessibility across various immune cell types in multi-task fashion, effectively decoding the regulatory syntax underlying immune cell differentiation. While ATAC-seq is commonly used to measure regional accessibility, it also provides high-resolution profiles, the distribution of Tn5 insertion sites, that offer additional insights into the precise location and strength of TF binding sites. Here we demonstrate that modeling ATAC-seq profiles alongside accessibility consistently improves predictions of differential chromatin accessibility across cell types. Moreover, we also find that multi-task learning across related immune cell types consistently outperforms single-task models. To understand what additional information bpAI-TAC learns from ATAC-seq profiles, we systematically compare sequence attributions from models trained with and without ATAC-seq profiles. We identify novel motifs with strong effect sizes that emerge only when profile data is included. Our findings suggest that modeling ATAC-seq at base-pair resolution enables the model to learn a more nuanced and sensitive representation of the cis-regulatory syntax driving immune cell-specific chromatin landscapes.

## Introduction

The formation of distinct cell types is driven by differential gene expression, which is tightly regulated by thousands of transcription factors (TFs) that bind to open chromatin regions (OCRs) and recruit the transcriptional machinery [1]. TFs recognize specific DNA sequence motifs, and their combinatorial interactions encode a complex regulatory syntax embedded within cis-regulatory elements (CREs) throughout the genome [2, 1]. This cis-regulatory syntax orchestrates the spatial and temporal control of gene expression in a cell type–specific manner. Chromatin accessibility, an indicator of CRE activity, can be measured genome-wide using ATAC-seq [3], which leverages the Tn5 transposase to cut and insert sequencing adapters at accessible regions of the genome. The number of Tn5 insertions serves as a proxy for regional accessibility [4], but the signal is typically aggregated over hundreds of base pairs, making it difficult to pinpoint the specific bases bound by TFs that drive accessibility changes [5].

To decode the regulatory syntax underlying chromatin accessibility, deep sequence-to-function (S2F) models have been developed to learn the relationship between DNA sequence and observed chromatin signals from genome-wide measurements [6, 7, 8, 9]. Previously, we developed AI-TAC, a multi-task S2F model that accurately predicts chromatin accessibility across diverse mouse immune cell types from DNA sequence [7]. While AI-TAC successfully learns accessibility-associated motifs, recent studies demonstrate that the fine-scale distribution of Tn5 insertions contains additional information about the location and binding strength of TFs in OCRs in the so-called “TF footprints” [10, 11, 12]. Models such as BPnet [13] and its ATAC-seq successor ChromBPnet [14] have successfully demonstrated that training on these base-pair (bp) resolution profiles enhances discovery of TF motifs and enables investigation of footprints that determine TF binding strength [8, 9]. Yet, it remains unclear if base-pair modeling also improves the models’ ability to capture the regulatory syntax underlying differences between cell-type specific accessibility.

Here, we present bpAI-TAC, a multi-task S2F model that predicts ATAC-seq counts at base-pair resolution across 90 immune cell types (Figure 1a) [4]. In contrast to ChromBPNet, which trains one model per cell type, bpAI-TAC uses a single multi-task model to enable direct comparisons across cell types, reduce computational burden, and ensure consistency across predictions. We show that incorporating base-resolution ATAC-seq profiles improves the model’s accuracy in predicting differential chromatin accessibility and enables high-resolution decoding of regulatory grammar. We also evaluate the impact of training resolution, model architecture, and objective functions on performance, and find that multi-task learning consistently improves prediction performance without requiring explicit bias factorization. Finally, by analyzing sequence attributions from bpAI-TAC and comparing them to those from AI-TAC, we identify additional TF motifs and candidate regulatory elements that are only uncovered when the model learns from base-pair resolution profile data.

**Fig. 1.**
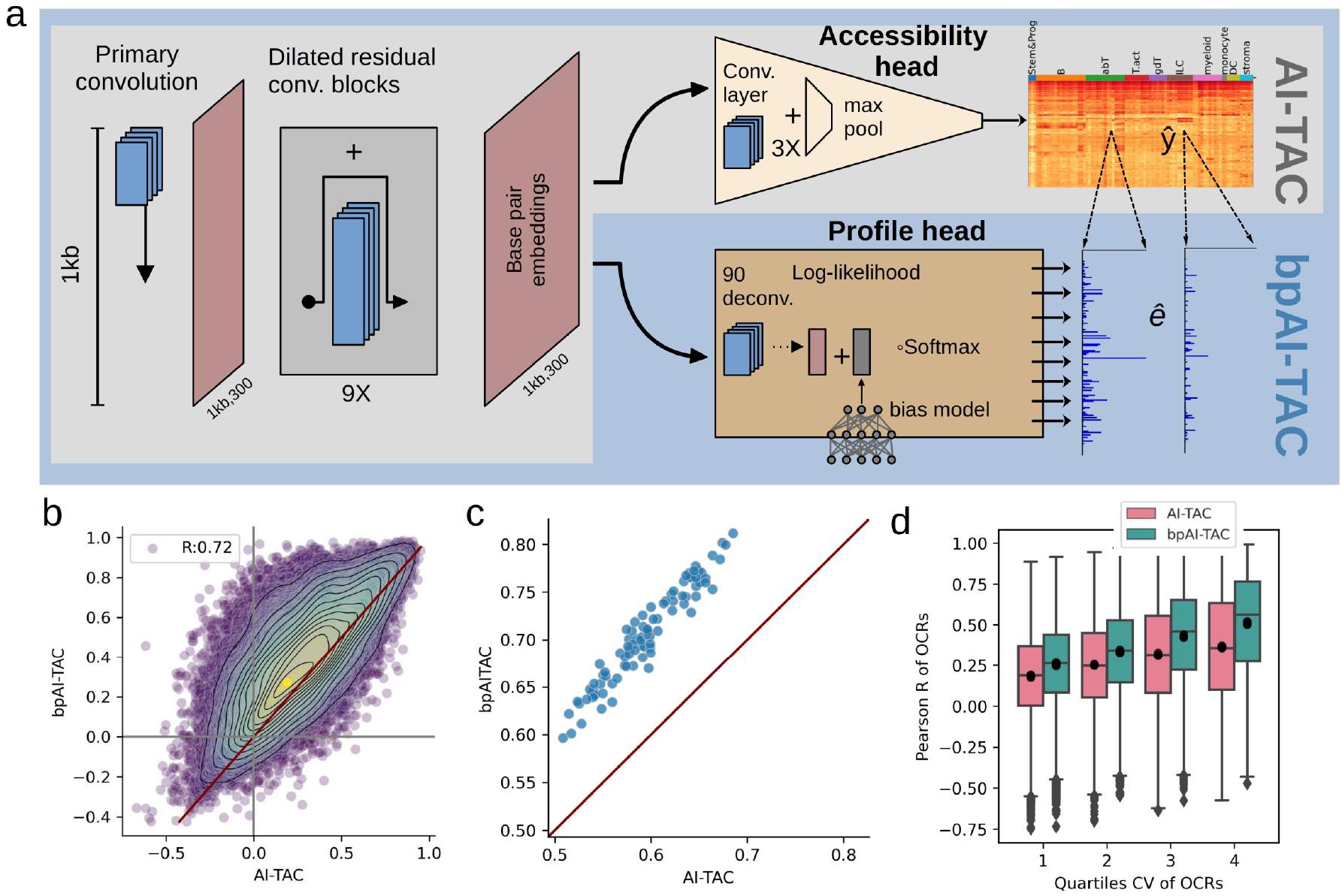
BpAI-TAC enhances performance via dual-head learning. a) BpAI-TAC is a multi-task CNN trained to predict base-pair resolution Tn5 insertions across 90 cell types. Its dual-head architecture jointly predicts chromatin accessibility (total Tn5 insertions within ±125 bp of peak centers) and Tn5 insertion profiles, while correcting for Tn5 sequence bias. b) Pearson correlation comparison across cell types between BpAI-TAC and AI-TAC for chromatin accessibility predictions on held-out OCRs. c) Pearson correlation (R) across cell types on held-out OCRs for both models. d) Boxplots of OCR R values grouped by quartiles of coefficient of variation across cell types.

## Results

### BpAI-TAC leverages ATAC-profiles to improve predictions of differential chromatin accessibility

AI-TAC is a convolutional neural network (CNN) designed to predict chromatin accessibility of OCRs across mouse immune cell types [7]. Unlike AI-TAC, bpAI-TAC predicts base-pair resolution Tn5 insertion counts (Figure 1a). Following previous model architectures [14, 13], bpAI-TAC takes as input 1kb genomic DNA sequence and produces two outputs: the “profile head” predicts the probability of Tn5 insertions for each of the central 250 bp, and the “accessibility head” predicts the aggregate number of insertions for the entire region. These outputs can be combined by multiplying the predicted profile with the predicted accessibility to reconstruct base-wise Tn5 insertion counts [13, 8] (Methods).

While accurately modeling bp-profiles is important to derive clean footprints from the model (i.e. inferring the exact binding location and strength of a TF) [9, 14], our analysis primarily focuses on how well these models can predict differences in chromatin accessibility between cell types. To quantify model performance, we primarily use Pearson correlation R of predicted versus observed accessibility of held-out OCRs across cell types as a metric to compare bpAI-TAC to an equivalent version of AI-TAC (Figure 1b,c). The equivalent version of AI-TAC uses the exact same architecture as bpAI-TAC, but without training on the bp-profile predictions. bpAI-TAC consistently outperforms AI-TAC in predicting differential accessibility (Figure 1b, S1a), suggesting that it learns additional grammar relevant for cell type-specific chromatin accessibility from the provided profiles. Furthermore, bpAI-TAC also improves predictions across test set OCRs for all cell types (Figure 1c, S1b), suggesting that the profile information also helps the model to discover more cell type specific factors that contribute to the variance between OCRs. When stratified by the coefficient of variation (CV) across cell types, we observe the largest performance gains in regions with high variability (Figure 1d), highlighting bpAI-TAC’s enhanced ability to learn motifs associated with dynamic, cell type–specific regulation. In summary, incorporating ATAC-seq profiles into model training substantially enhances the model’s ability to predict chromatin accessibility differences across cell types, indicating a more refined understanding of cis-regulatory grammar.

### Tn5 profiles improve open chromatin predictions consistently across of modeling choices

BpAI-TAC can be trained using any objective function that measures differences between the prediction of the number of Tn5 insertions per base pair, computed as the product of the profile and accessibility heads. To determine the optimal modeling objective, we evaluated regular and composite loss functions, which weight the errors of the profile and accessibility heads separately using a hyperparameter (*λ*), as in (Chrom)BPNet [13, 14] (Figure 2a, Methods). While composite losses offer flexibility, they require careful tuning of *λ* on validation data (Figure S2).

**Fig. 2.**
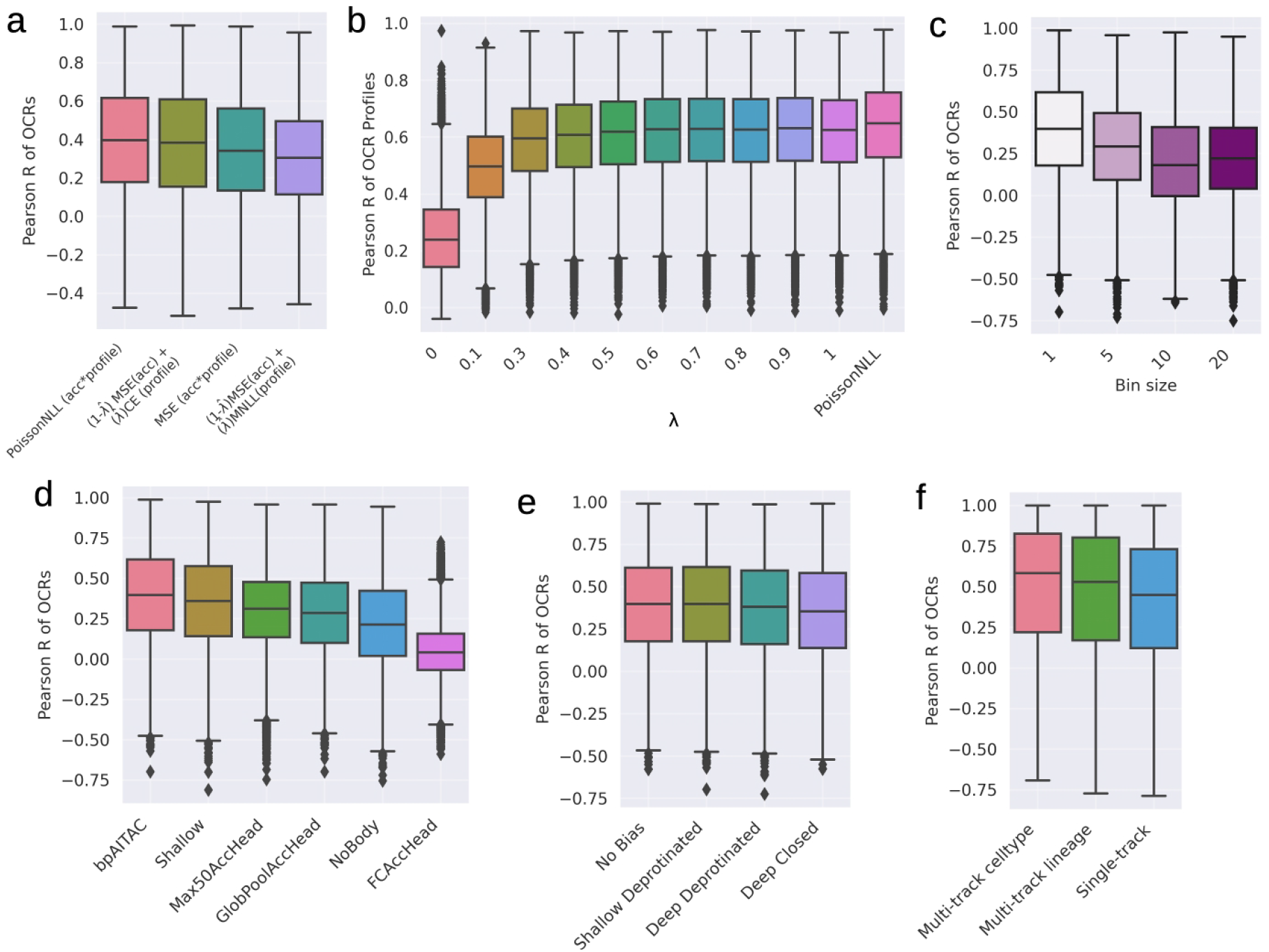
Tn5 profiles consistently enhance open chromatin prediction across bpAI-TAC modeling strategies. a) Distribution of Pearson correlation coefficients (R) for open chromatin regions (OCRs) across cell types, comparing four different loss functions. b) Pearson R distributions for profile predictions from models trained with composite loss functions combining mean squared error (MSE) and negative multinomial loss. The contribution of profile loss was modulated by the profile loss fraction (*λ*). c) Pearson R distributions for OCR predictions from bpAI-TAC models trained on Tn5 profiles at varying resolutions. Lower-resolution profiles were generated by binning raw Tn5 signal into windows of size 5, 10, or 20 base pairs. d) Ablation experiments on the bpAI-TAC architecture, including variations with a reduced model body (Shallow, NoBody) and alternative designs for the accessibility prediction head (GlobPoolAccHead, Max50AccHead, FCAccHead). e) Comparison of four Tn5 bias correction strategies and their impact on OCR prediction. The strategies include: (1) no bias correction, (2) a shallow CNN trained on protein-free DNA, (3) a deep CNN trained on protein-free DNA, and (4) a deep CNN trained on aggregated closed OCR regions. f) Pearson R distributions across cell lineages for three modeling approaches: (1) a Multi-Track Celltype model trained to predict accessibility in 90 individual cell types and aggregated into 10 lineages, (2) a Multi-Track Lineage model trained directly on 10 lineages, and (3) a Single-Track model ensemble, with one model trained per lineage.

First, we compared different loss functions, composite and regular, on how well they model differences of OCRs between cell types (Pearson R OCRs) and differences between OCRs within cell types (Pearson R cell types). Consistent with the theoretical distribution of sequencing reads, we found that training our model on the Poisson negative log-likelihood (PNLL) loss returns the best results (Figure 2a, S3a). The mean squared error (MSE) loss on the number of Tn5 insertions performs slightly worse in both directions. On the other hand, the composite loss approximation of the PNLL loss as MSE of logged counts and the multinomial negative log-likelihood (MNLL) [13] performed consistently worse than both regular per-base count losses. While the composite loss with the MNLL represents the theoretically correct approximation of the Poisson loss [13], we observe that a mixture of MSE and cross-entropy (CE) loss achieves better performance for predicting differences between cell types, but lags behind for differences between OCRs (Figure S3a), possibly because CE, unlike MNLL, does not focus high-count regions (Methods).

To understand the influence of the composite losses’ hyperparameter lambda on the model’s predictions, we trained models with different ratios between profile and accessibility head from 0 (only training on accessibility head, i.e. AI-TAC) to 1 (only training on profile head). As expected, the prediction of profiles consistently improves until the model is only trained on the CE loss (Figure 2a, S2a,b,c). Interestingly, also for the profiles, the PNLL performs better than the composite loss, suggesting that the loss better captures the distribution of the Tn5 insertions. We found that models which included base resolution profiles in their objective improved overall accessibility prediction compared to AI-TAC, even without tuning (Figure S2c, d). However, when model training was focused too much towards profiles (lambda ¿ 0.6) the performance for predictions of total chromatin accessibility across cell types declined again and became random for a model that was only trained on profiles (lambda = 1.0).

To investigate if multi-tasking helps our model to extract more information from the data than an ensemble of single-track models, we compared a lineage-averaged multi-task model to an ensemble of 10 lineage-specific single-task models (10 lineages in 90 cell types), and to the lineage-averaged predictions from our model trained on individual cell types. Both models trained as multi-task models show improved predictions across cell lineages compared to the ensemble of single-task models (Figure 2c). Additionally, predictions are also improved in the other direction for individual cell lineages across held-out OCRs, suggesting that the multi-task model does not only improve scaling between tasks, but also extracts more information about the regulatory code (Figure S3b).

Next, we evaluated architectural choices (Figure 2d, S3c). First, we replaced the deep architecture that used 9 dilated residual convolutional blocks with a shallower version that only used 4 blocks (Shallow), and observed minimal impact on performance. When we replaced the body of the model entirely with a single convolutional layer (NoBody), we observed significantly decreased performance, likely due the inability of this model to learn positional effects of motifs, motif context, or motif interactions. Replacing the entire accessibility head with global mean-pooling of the base pair representations over the entire OCR (GlobPoolAccHead), as used by (Chrom)BPnet, also had a negative impact on the model performance. Replacing the global mean-pooling with smaller max-pooling over 50 base pairs (Max50AccHead) led to even worse performance. On the other hand, when we replaced the scalar head with a massive fully connected layer (FCAccHead), instead of multiple convolutions and pooling layers, the model significantly underperformed and reached almost random performance, reinforcing that convolutional structures are better suited for decoding regulatory syntax.

Then, we investigated methods to account for Tn5-enzyme’s sequence bias on bpAITAC’s ability to learn differential accessibility (Figure 2e, S3d). As previously shown [14], the Tn5-enzyme possesses a strong sequence preference that impacts the local distribution of Tn5 insertions in OCRs [11, 10]. This strong sequence bias distorts the distribution of Tn5 insertion sites around TF binding locations and can hide TF footprints. Without correction for this sequence bias, a S2A model may have to learn the sequence motifs that are associated with the Tn5 enzyme and therefore might miss signals from actual TF binding sites [13, 14, 11]. To investigate the impact of Tn5 sequence bias, we followed the strategy of (Chrom)BPnet and modeled the distribution of Tn5 insertions separately from the accessibility to add the log-likelihood from a bias model before the softmax function (Methods, Figure 1a). We trained three bias models using the bpAI-TAC architecture, optimizing only the profile loss: (1) a model trained on protein-free DNA from [10]; (2) a model trained on aggregated profiles from closed OCRs across all 90 cell types; and (3) a shallower variant trained on the same protein-free data. Closed OCRs were defined using peak-calling probabilities (Methods). Since individual closed regions contain few reads, we aggregated counts across cell types where the region was consistently closed. This strategy not only increases signal but also reduces training time. All three models produced highly concordant bias profiles (Figure S4a,b). When evaluated on held-out chromosomes and protein-free DNA, both deep models generalized well, accurately predicting each other’s profiles (Figure S4c,d). We also trained a model without any bias correction to assess its necessity. All four models achieved similar predictive performance overall (Figure 2e). Notably, while the protein-free models slightly underperformed in cross-cell type predictions across OCRs (Figure S3d), the model without bias performed comparably—or even slightly better—when predicting differences between cell types and OCRs (Figure 2d, S3d). These results suggest that, despite the known interpretability advantages of bias modeling [14] sufficiently expressive models like bpAI-TAC can effectively learn both biological and assay-specific signals directly from the data.

One of the primary bottlenecks in modeling chromatin accessibility at base-pair (bp) resolution using a multi-task framework is the substantial memory requirement. Representing every nucleotide dramatically increases data size—by a factor proportional to sequence length. However, since transcription factor (TF) binding sites typically span only 5–20 bp [17, 18, 1], full bp-resolution may not be necessary to capture the key features of TF footprints. To test this hypothesis, we trained models using coarser profile resolutions of 5, 10, and 20 bp. In each case, we incorporated the Tn5 bias into the base-resolution log-likelihood, then aggregated predictions within each bin before applying the softmax (Methods). As with the base-resolution model, we evaluated predictions against measured accessibility on a held-out test set. Our results indicate that reduced resolution models consistently underperform compared to the single base pair model, yielding lower accuracy in predicting total chromatin accessibility (Figure 2f, S3e). We hypothesize that this performance drop arises from coarse binning, which may obscure fine-scale TF footprints, either by including only parts of a footprint or by merging signals from different TFs, particularly those without strong, multi-base-pair valleys.

### BpAI-TAC learns additional motif syntax that drives immune cell differentiation

Biologically meaningful improvements in prediction would require bpAI-TAC to learn a more comprehensive cis-regulatory syntax. To explore this, we conducted careful model interpretation, comparing motifs learned by bpAI-TAC with those of AI-TAC at OCR basis. We identified a set of 1,082 sequences that bpAI-TAC consistently predicted well, and better than AI-TAC, across five model initializations (Figure 3a, Methods). To each of these sequences, we applied DeepLIFT and identified the motifs contributing to predictions across 90 cell types [19]. Visual inspection showed that bpAI-TAC consistently identified clear motifs in these sequences, while AI-TAC either missed them entirely or only partially (Figure 3b). To systematically determine motif clusters that are missed most frequently by AI-TAC, we extracted motif seqlets from attribution maps of both models, jointly clustered them, and identified motifs recognized by either or both models (Methods). We found 23 motif clusters that were significantly enriched in bpAI-TAC attributions compared to AI-TAC (Fisher exact test, Benjamini-Hochberg correction, FDR *<* 0.05, Figure 3c).

**Fig. 3.**
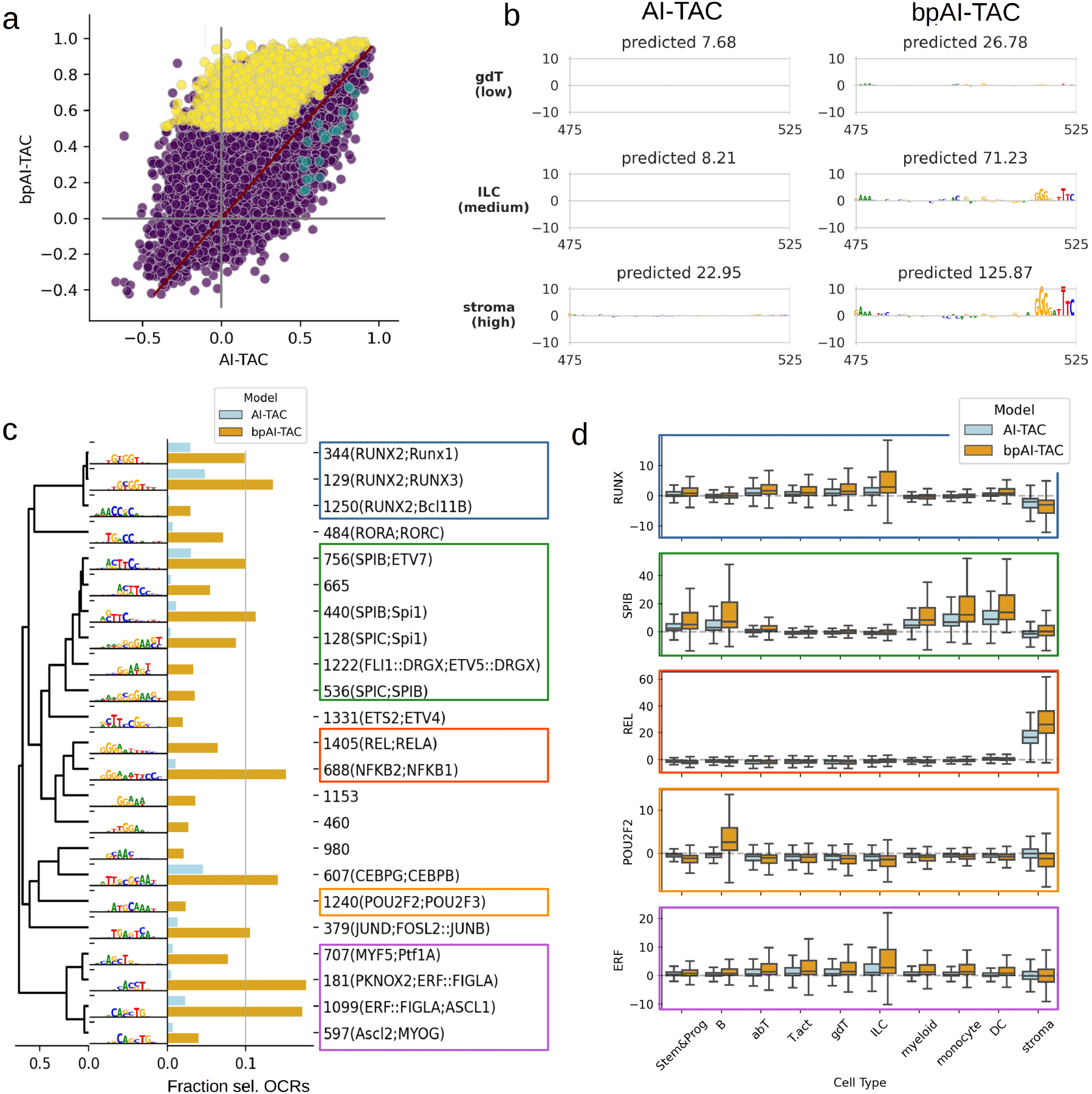
BpAI-TAC learns additional motifs that drive immune cell differentiation from TF footprints. a) Comparison of Pearson correlation R of OCRs across cell types as in Fig 1a), with selected regions for attribution analysis in yellow. b) An example of sequence attributions from AI-TAC and bpAI-TAC for one of the selected yellow regions in a). Attributions across three different cell lineages are shown, increasing in experimentally measured accessibility from top to bottom. c) Combined motifs from seqlet clustering sorted on a tree with agglomerative clustering and average linkage on their correlation distance. Seqlets were extracted from sequence attributions across cell lineages and hierarchically clustered (Methods). Bar plots behind motifs show the fraction of all sequences in which the motif cluster was found in attributions of any lineage from AI-TAC (light blue), and attributions from bpAI-TAC (golenrod). Cluster identifiers are shown behind with associated transcription factor names in brackets (TomTom, q-value *<* 0.05, Jaspar non-redundant vertebrate database [15, 16]) d) The difference in accessibility predictions on 4000 dinucleotide shuffled sequences with and without motifs inserted. The difference in bpAI-TAC’s predictions are displayed in orange, and the difference in AI-TAC’s predictions are in blue.

The three most enriched motif cluster groups were associated with RUNX-like (cluster 344, 129, 1250), SPIB-like (cluster 756, 665, 440, 128, 1222, 536), REL-like (Cluster 1405, 688), POU2F-like (cluster 1240), and ERF-like motifs (cluster 707, 181, 1099, 597). To determine if AI-TAC misses these motifs globally or just in the context of these sequences, we inserted the motifs into 4,000 randomly shuffled sequences and assessed the predicted effect sizes of each model (Figure 3d). While these motifs also influenced AI-TAC predictions across cell types, the effect sizes differed significantly across many cell lineages (Wilcoxon rank-sum test). bpAI-TAC predicts much larger effect sizes of POU2F2-like motifs in B-cell, REL-like motifs in stroma, of RUNX-like motifs in ILC, and T-cells, and SPIB-like motif in stem, B, myeloid, monocyte and DC cell lineages. These results suggest that bpAI-TAC does not discover entirely new motifs, but improves the estimation of their effect sizes in different contexts.

We hypothesized that bpAI-TAC’s improved performance arises from its ability to learn transcription factor (TF) footprints directly from base-pair–resolution chromatin profiles. To test this, we generated predicted TF footprints (see Methods), setting the Tn5 bias to zero during inference to return solely the residual signal, representing what the model has learned about TF binding patterns [14]. When we inserted motif groups into randomly shuffled background sequences, bpAI-TAC produced distinct footprint-like patterns at motif locations for all four motif groups, in contrast to AI-TAC, which exhibited increased noise at these positions (Figure S5). These results suggest that base-pair–level profiles help bpAI-TAC learn the precise functional impact of motifs, consistent with the findings from our ablation experiments (Figure S2d,e). In contrast, AI-TAC may recognize motif presence but fails to capture their regulatory effect size in context, resulting in substantially smaller predictive impacts in relevant cell types.

## Discussion

We demonstrated that modeling ATAC-seq data at bp-resolution not only improves interpretability of the models’ predictions [14, 9, 8], but also significantly improves the ability to capture differential regulatory activity across closely related cell types. Our model, bpAI-TAC builds on modeling choices of (Chrom)BPnet [13, 14], but focuses on modeling the data in a comparative multi-task fashion, which facilitates training and improves comparisons of differential activity across cell types.

Moreover, our results highlight that model performance is highly sensitive to architectural and loss function choices. In contrast, the choice of the Tn5 bias model, though important for interpretability of footprints [14], had relatively little effect on overall predictive performance. Models trained without explicit bias correction still learned meaningful motif syntax from the profile data, leading to improved performance. We hypothesize that bias correction is less critical in the dual-head framework because we decouple profile and accessibility predictions. This separation enables accessibility attributions to selectively ignore sequence features, such as the Tn5 motif bias, that primarily affect profile shape but are unrelated to chromatin accessiblity (Figure S6).

When we investigated differences in the learned sequence grammar to AI-TAC, we observed that AI-TAC systematically overlooked or underestimated the effect size of various motif groups in certain cell-type specific contexts. While bpAI-TAC showed improved alignment of residual profiles (after Tn5 bias removal) with known motif positions, the resulting deep footprints were less pronounced than those observed in ChromBPnet [14] (Figure S5). For several motifs, we did not detect clear multi-base-pair valleys typically associated with high-confidence footprints; instead, we observed narrower, spike-like patterns with reduced variance. These may reflect weakly bound TFs or result from suboptimal bias correction used here, similar to observations in previous studies [14]. It remains unclear whether this is due to our multi-task architecture, limited sequencing depth (2.5–40 million reads per cell type, 15 million on average), the quality of the protein-free or aggregated data used for bias modeling, or other design choices.

In summary, we present a multi-task convolutional framework for modeling ATAC-seq data at base-pair resolution. Consistent with prior studies [13, 14, 9], we find that incorporating base-resolution information significantly enhances both predictive performance and interpretability. While previous approaches have emphasized the model’s ability to create footprint with explicit bias factorization [13, 14], the dual-head design, jointly modeling total accessibility and Tn5 insertion profiles, proves effective without requiring separate bias modeling. Additionally, we demonstrate that multi-task learning consistently improves predictive accuracy across cell types, highlighting its value for comparative regulatory modeling.

## Methods

### Data processing

All models were trained using ATAC-seq data from 90 mouse immune cell types collected by the Immunological Genome Project [4]. As input to the models, we used one-hot encoded sequences of length 998 bp around ATAC-seq peaks from the mm10 mouse genome. Open chromatin regions (peaks) were previously called in [4]. Roughly, 2 to 181 samples were grouped with hierarchical clustering on various cut-offs to estimate the peak summits for cell type and lineages. This resulted in 518,845 ATAC-seq OCRs. We divided chromatin regions into training, validation, and test sets, leaving out chromosomes 12 and 15 for validation during training and 11 and 16 for testing model performance. Cumulative ATAC-seq counts were calculated by summing up the number of raw Tn5 insertions in the 250 bp region surrounding the ATAC-seq peak location. We used quantile normalization for these sums to account for different sequencing depths of 2.5-40 million fragments. The ATAC-seq profiles were unaffected by this normalization.

### BpAI-TAC model

BpAI-TAC is composed of a body of convolutional layers and two output heads, one that predicts the sum of Tn5 insertions (accessibility head), and one that predicts the ATAC-seq profile (profile head). The body of the model starts with a convolution of width 25, followed by 9 dilated convolutions of width 3 with residual skip connections wherein dilations start at a size of 2 and double each layer. All convolutions in the body of the model have 300 filters and are followed by ReLU activation. The model then branches into two separate heads. The profile head uses a single convolution of width 25 with 90 filters, one for each cell type. If used, we then add the predicted log-likelihood of the Tn5 sequence bias model for the associated input sequence (i.e. values from bias models before Softmax), and apply a Softmax function to transform the log-likelihoods into a probability distribution of Tn5 insertions for each base in the 250 bp window. The scalar head consists of three repetitions of max pooling with width 5 followed by a convolution of width 3 with 300 filters and a ReLU activation. The scalar head ends with a fully connected layer, which returns predictions for 90 cell types simultaneously.

### Bias-model training

All Tn5 bias models were only trained in predicting profiles correctly, without scalar head loss, using only the JSD profile loss. We trained three Tn5 bias models with the bpAI-TAC architecture: 1) protein-free DNA from [10], and 2) cumulative profiles from closed OCRs. The third bias model used the shallow bpAI-TAC architecture and was trained on protein-free DNA profiles. Protein-free DNA had been generated for 25 selected chromatin regions based on overlap with a manually selected set of key transcription factors and differentiation-related genes [10]. To train a bias model on closed OCR profiles from the 90 cell types, we identified closed OCRs using the MACS2 peak calling probability given the number of insertions in each cell type individually (*p >* 0.25). Individual closed regions contain only very few reads in the cell types in which they are inaccessible. To generate profiles for closed regions with larger number of insertions, and to reduce the number of data points used in this training strategy, we used the aggregated number of Tn5 insertions across cell types in which the OCR is closed.

### Model ablations

#### Binning ablation

To investigate whether lower resolution accessibility profiles could also improve predictions of chromatin accessibility, we binned the ATAC profiles in windows of 5, 10, and 20 base pairs. In the profile head, we sum across the sequence within bins prior to taking the Softmax. Thus, the predicted Tn5 bias is also included in this binning, and can be accounted for with a bias model. This ensures fair comparison with the no-binning condition.

#### Architecture ablations

To investigate how model architecture choices influence the performance of bpAI-TAC, we investigated changes in the sequence embedding body of the model, and the two prediction heads. First, we reduced the number of dilated convolutional layers in the body from 9 to 4 (Shallow) and also completely removed the dilated convolutions (NoBody). For the accessibility head, we replaced the repeated convolutional layers with poolings by a single fully connected layer (FCAccHead), a single global average pooling strategy (GlobPoolAccHead), and a pooling at a large window max of 50bp (Max50AccHead). For all these changes, the other modeling choices remained the same as in bpAI-TAC.

#### Loss ablations

We compared regular and composite loss functions. Regular loss functions measured the error of the number of Tn5 prediction per base pair directly. These predictions were explicitly computed by multiplying the predicted profile probabilities with the predicted total counts (accessibility) from the two bpAI-TAC output heads, and then compared to the measured count data. As regular losses, we tested Poisson Negative Log Likelihood (PNLL) loss and mean squared error (MSE). We also tested composite losses that contain a mixture parameter *λ* which represents the fractional weight of the profile loss compared to the total accessibility objective, consisting of MSE of logged counts:

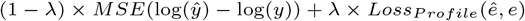

When *λ* = 1 the model is trained only on the profiles, while when *λ* = 0 the model is solely trained on the total accessibility counts, i.e. AI-TAC. We assessed the predictive power of two composite losses: First, using the multinomial negative log likelihood (MNLL) for the profiles, which in combination with MSE of logged counts approximates the PNLL [13]. Second, a composite loss of the MSE of logged counts and the cross entropy (CE) loss of the profiles.

### Sequence attribution and motif analysis

We selected a set of 1,082 sequences that were consistently well and better predicted with bpAI-TAC compared to AI-TAC (Supplementary Methods). We used DeepSHAP [19] to compute sequence attributions for 10 cell lineages by summing the predictions for the cell types in each lineage before applying DeepSHAP. We extracted motifs from the attribution maps (i.e. seqlets), clustered them, and used Tomtom [15] to assign TF names (details in Supplementary Methods). Finally, we determined motif clusters that were present significantly more often in bpAI-TAC than AI-TAC.

### Deep footprints and motif marginalizations

We used the combined motifs derived from the alignment of seqlets in the previous section for our marginalization analysis. We introduced the extracted and clustered motifs into 4000 randomly dinucleotide-shuffled OCR sequences from the test dataset and compared the predicted profiles and predictions from the bpAI-TAC model with predictions from AI-TAC. Additionally, we created footprints with the model’s profile predictions by setting the bias of the model to zero [14]. This strategy returns only the part of the profile that is unrelated to the Tn5 bias, and therefore contains the TF footprints. We plotted the predicted profiles’ median, the 25% and 75% percentile and compared them against the median and percentiles from randomly shuffled sequences without the motif.

## Data and code availability

This paper analyzes existing, publicly available data from Yoshida et al. 2019 DOI: 10.1016/j.cell.2018.12.036 [4]. The GEO accession number for the ATACseq data reported in this paper is GSE100738. Processed ATAC-seq data and called peaks can be found at: https://sharehost.hms.harvard.edu/immgen/ImmGenATAC18_AllOCRsInfo.csv. All original code has been deposited and is publicly available at https://github.com/nuriachandra/bpAITAC.

## Acknowledgments

This research was in part made possible by NIH-R24 (5R24AI072073-18), the Washington Research Foundation and the Carl-Zeiss-Stiftung. The funders had no role in study design, data collection and analysis, decision to publish or preparation of the manuscript.

## Author contributions

Conceptualization: N.A.C., A.S., S.M., Y.H., J.B. Methodology: N.A.C., A.S., S.M. ; Data curation: A.S., Y.H., J.B. Software: N.A.C., A.S.; Investigation: N.A.C, A.S.; Formal analysis: N.A.C., A.S.; Visualization: N.A.C. A.S.; Validation: N.A.C.; Writing: N.A.C., A.S., S.M.; Supervision: A.S. and S.M.

## Supplementary Methods

### Composite loss functions

We compared regular and composite loss functions. Regular loss functions measured the error of the number of per base pair Tn5 prediction directly. These predictions were explicitly computed by multiplying the predicted profile probabilities with the predicted total counts (accessibility) from the two bpAI-TAC output heads, and then compared to the measured count data. As regular losses, we tested Poisson Negative Log Likelihood (PNLL) loss and mean squared error (MSE). We also tested composite losses that contain a mixture parameter *λ* which represents the fractional weight of the profile loss compared to the total accessibility objective, consisting of MSE of logged counts:

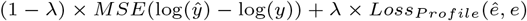

When *λ* = 1 the model is trained only on the profiles, while when *λ* = 0 the model is solely trained on the total accessibility counts, i.e. AI-TAC. We assessed the predictive power of two composite losses: First, using the multinomial negative log likelihood (MNLL) for the profiles. Second, a composite loss of the MSE of logged counts and the cross entropy (CE) loss of the profiles.

The composite loss of the MSE of logged counts and the multinomial negative log likelihood (MNLL) of the profiles approximates the PNLL loss Avsec et al. 2021 [13]:

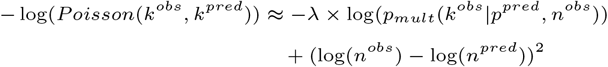

with

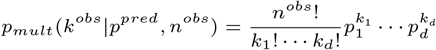

We also evaluated the composite loss of the MSE of logged counts and the cross entropy (CE). We did this because if we use the above relationship and substitute in *p*_*m*_*ult*, we can see that this composite loss represents a weighting of the cross entropy loss of each profile with the total counts of that region:

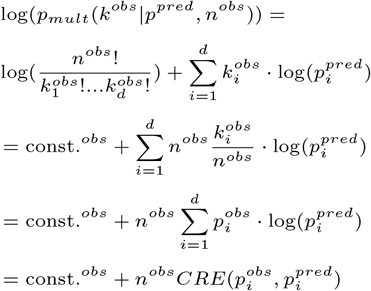

### Seqlet extraction & clustering

We used the following criteria to determine sequences that were consistently well and better predicted in bpAI-TAC compared to AI-TAC in five model initializations:

- Strong signal: *max*_*Celltype*_(accessibility) > 150.
- High variability across cell types: coefficient of variation > 1.
- Good prediction with bpAI-TAC: average Pearson correlation *R* > 0.5.
- Consistently better than AI-TAC: all ten initializations *R*_*bpAI−T AC*_ > all trials of*R*_*AI−T AC*_

For the resulting 1,081 sequences in the test set, We used DeepSHAP [19] to compute the attributions for the 10 cell lineages by summing the predictions for the cell types in each lineage before applying DeepSHAP. We extracted motifs from the attribution maps (i.e. seqlets), using rules about significance and number of subsequent base attributions. Hypothetical attribution scores (i.e. centered to zero at each position) for all four base-pairs were extracted for windows if it contained attributions for the present base that were above 1.96 of the standard deviation (equivalent to p-value *<* 0.05, two-tailed T-test) of attributions across all sequences, and the motif consisted of at least four of these significant bases allowing only for single insignificant base-pair gaps between significant positions. Seqlets were extracted from the attributions of bpAI-TAC and AI-TAC across all 10 lineages and then clustered with agglomerative clustering with complete linkage, using p-values of the strongest correlation of aligned motifs as a distance metric. Using p-values instead of correlations accounts for the length of the motif alignment. Clusters were assigned to motifs that shared at least a 0.05 p-value for their correlation with each other (Figure 3c).

## Supplementary Figures

**Fig. S1.**
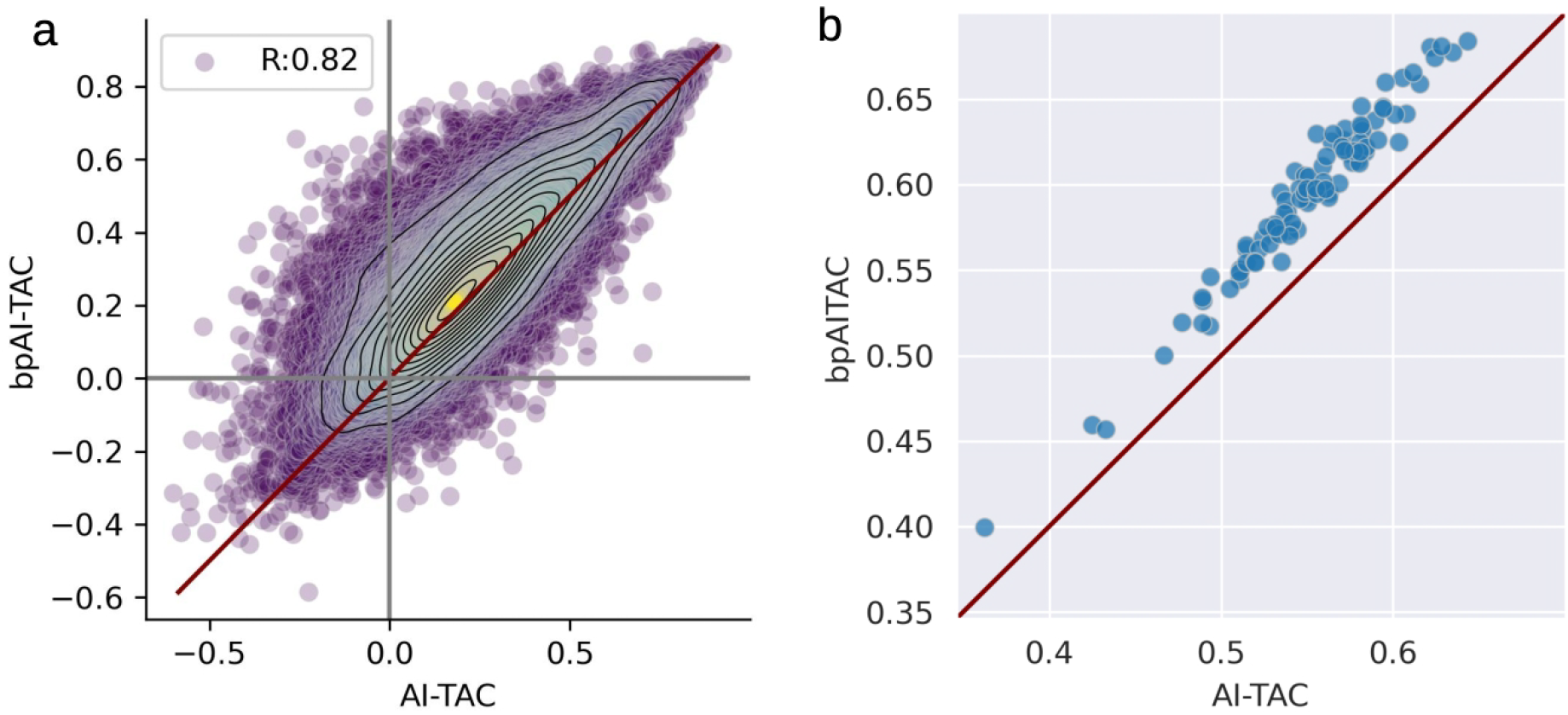
a) Spearman correlation of OCRs across cell types for bpAI-TAC and AI-TAC accessibility prediction. b) Spearman correlation of cell types across OCRs for bpAI-TAC and AI-TAC accessibility prediction.

**Fig. S2.**
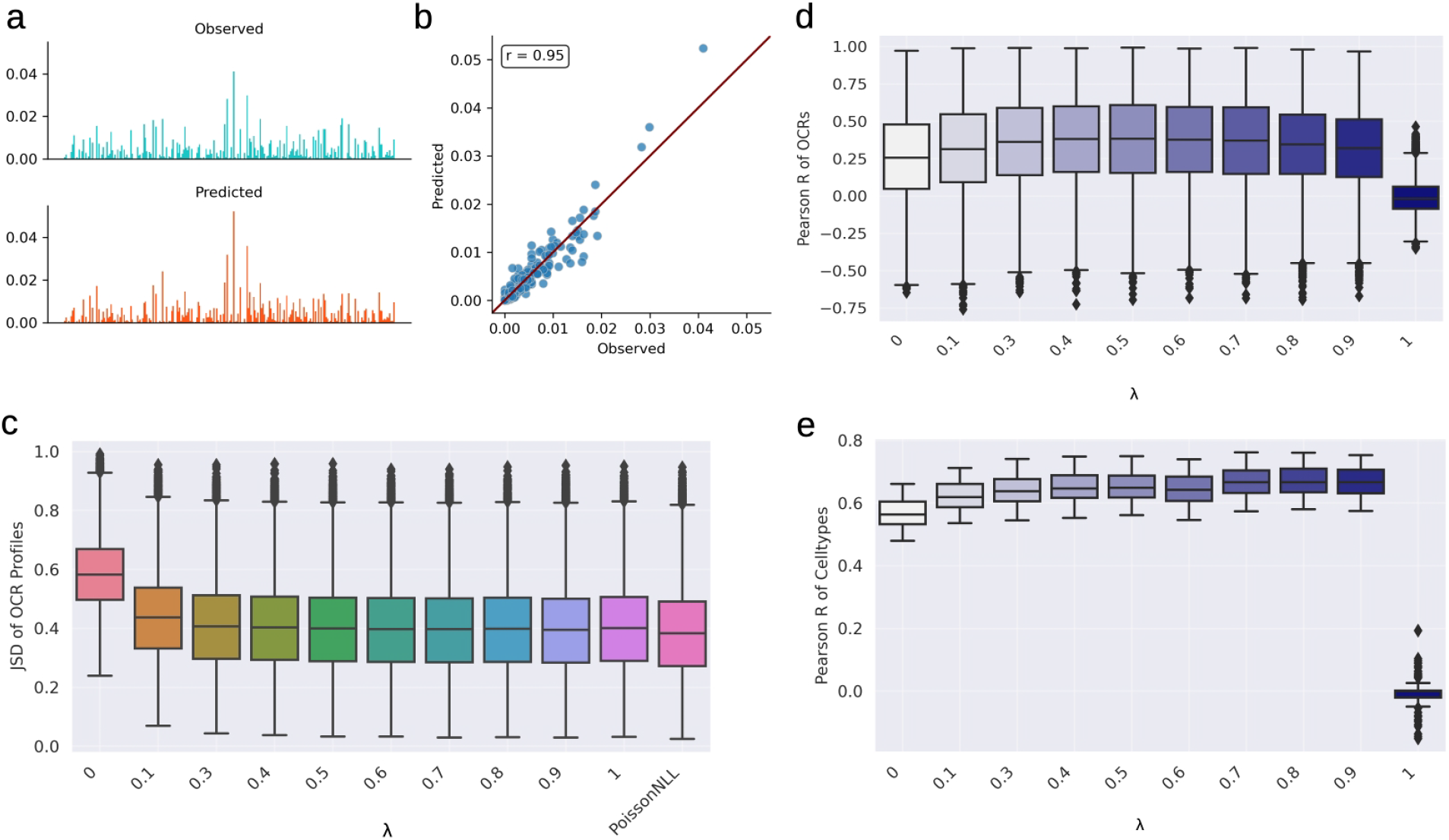
Optimization of the accessibility-profile ratio lambda of the composite loss function (MSE+CE) a) Predicted Observed profile in single OCR in a cell type b) Scatter plot between the predicted per base probabilities in a). c) Jensen-Shannon divergence (JSD) distribution between predicted and observed Tn5 insertion profiles. d) Pearson R of OCRs for different lambda ratios. e) Pearson R of Cell types for different lambda ratios

**Fig. S3.**
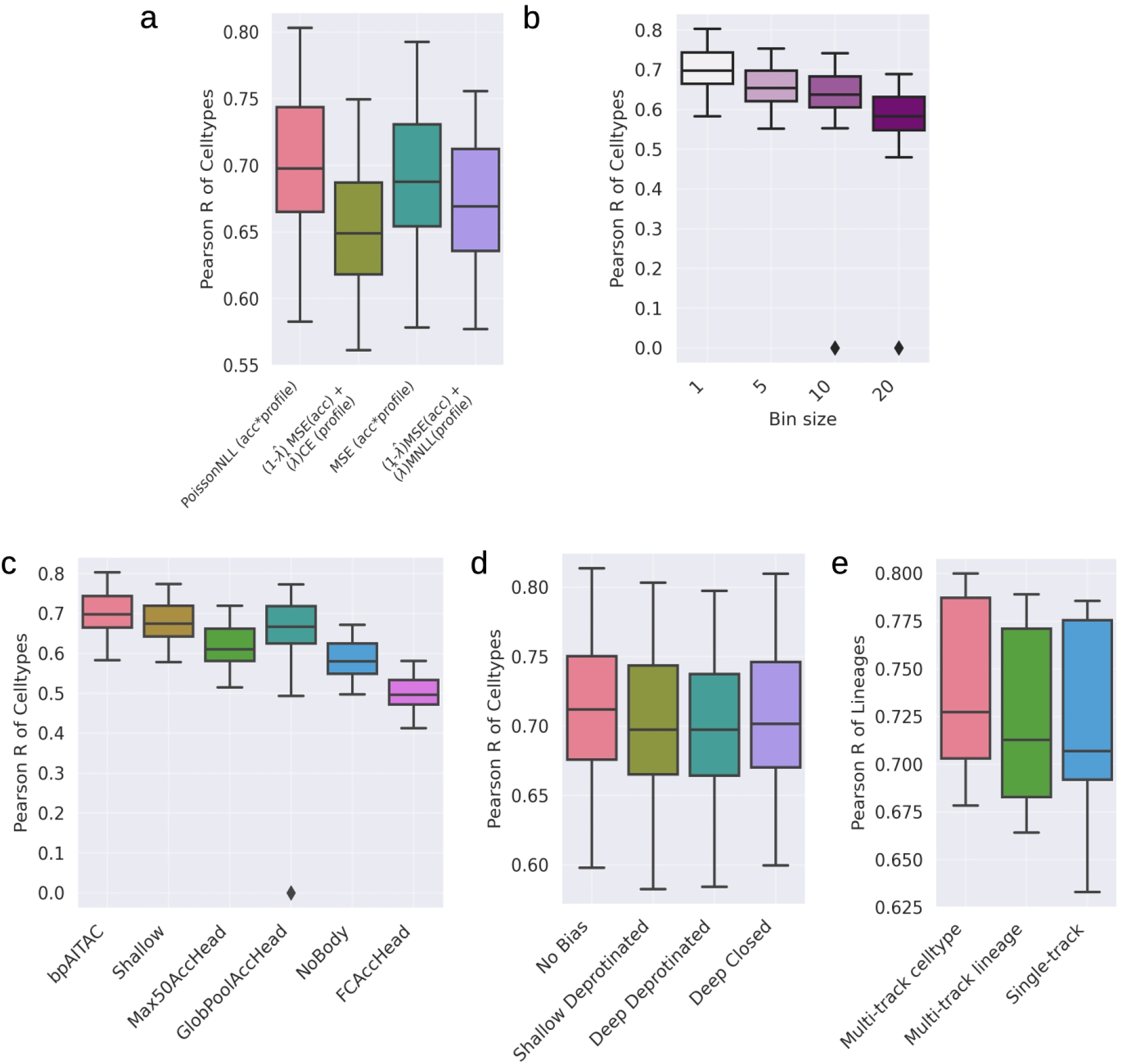
Predictions across test set OCRs for different cell types. a) Pearson R of cell types across OCRs for four loss functions. b) Pearson R of cell types across OCRs for bpAI-TAC trained on different resolution profiles. Profiles were binned (summed over regions of sizes 5, 10, and 20) to create lower resolution Tn5 profiles. c) Pearson R of cell types for BpAI-TAC architecture ablations, including reducing the size of the body (Shallow and NoBody), and modifying the accessibility head (GlobPoolAccHead, Max50AccHead, FCAccHead). d) Pearson R of cell types across OCRs for bpAI-TAC trained using four different Tn5 bias prediction approaches. From left to right these approaches are 1) no Tn5 bias prediction added, 2) a shallow CNN trained on protein-free DNA, 3) a deep CNN trained on protein-free DNA, and 4) a deep CNN trained on closed OCR regions. e) Pearson R of cell lineages across OCRs. The Multi-Track Celltype model is trained to predict accessibility in 90 different cell types, with predictions averaged into 10 different lineages. The Multi-Track Lineage model is trained to predict accessibility across 10 different lineages. The Single-Track model is an ensemble of the results from 10 individual models trained on individual lineages. U

**Fig. S4.**
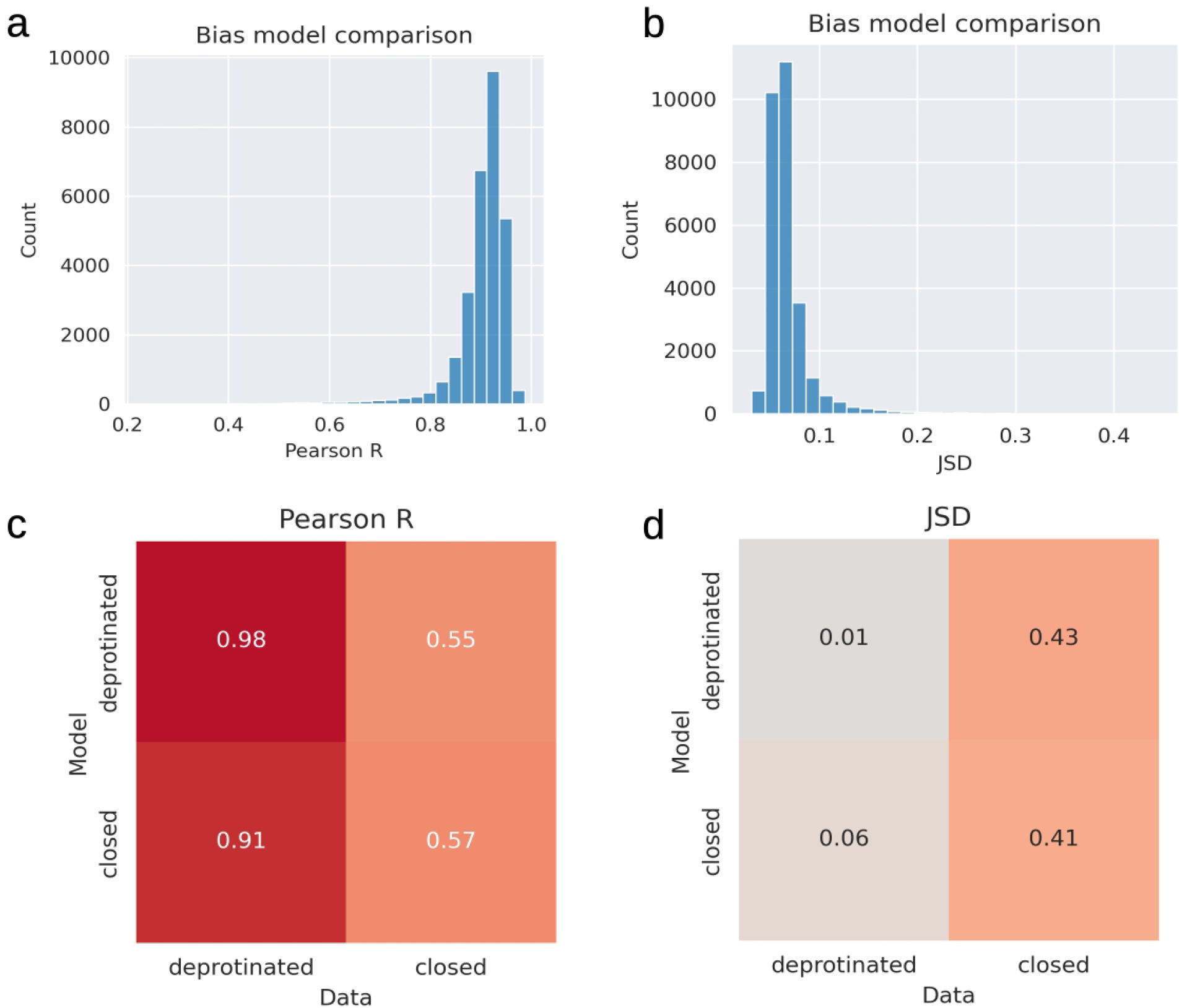
Comparison of Tn5 bias prediction models trained on aggregated closed DNA regions and protein-free DNA. All analyses are performed on regions of DNA from held-out chromosomes. a) Mean Pearson R across profile predictions from two models trained on protein-free and aggregated closed regions on held-out test regions of closed regions of DNA. b) Jensen-Shannon divergence (JSD) between protein-free and closed bias model predictions on held-out test regions of closed regions of DNA. c) Pearson R of observed and predicted closed and protein-free DNA for both types of bias model. d) JSD of observed and predicted closed and protein-free DNA for both types of bias model.

**Fig. S5.**
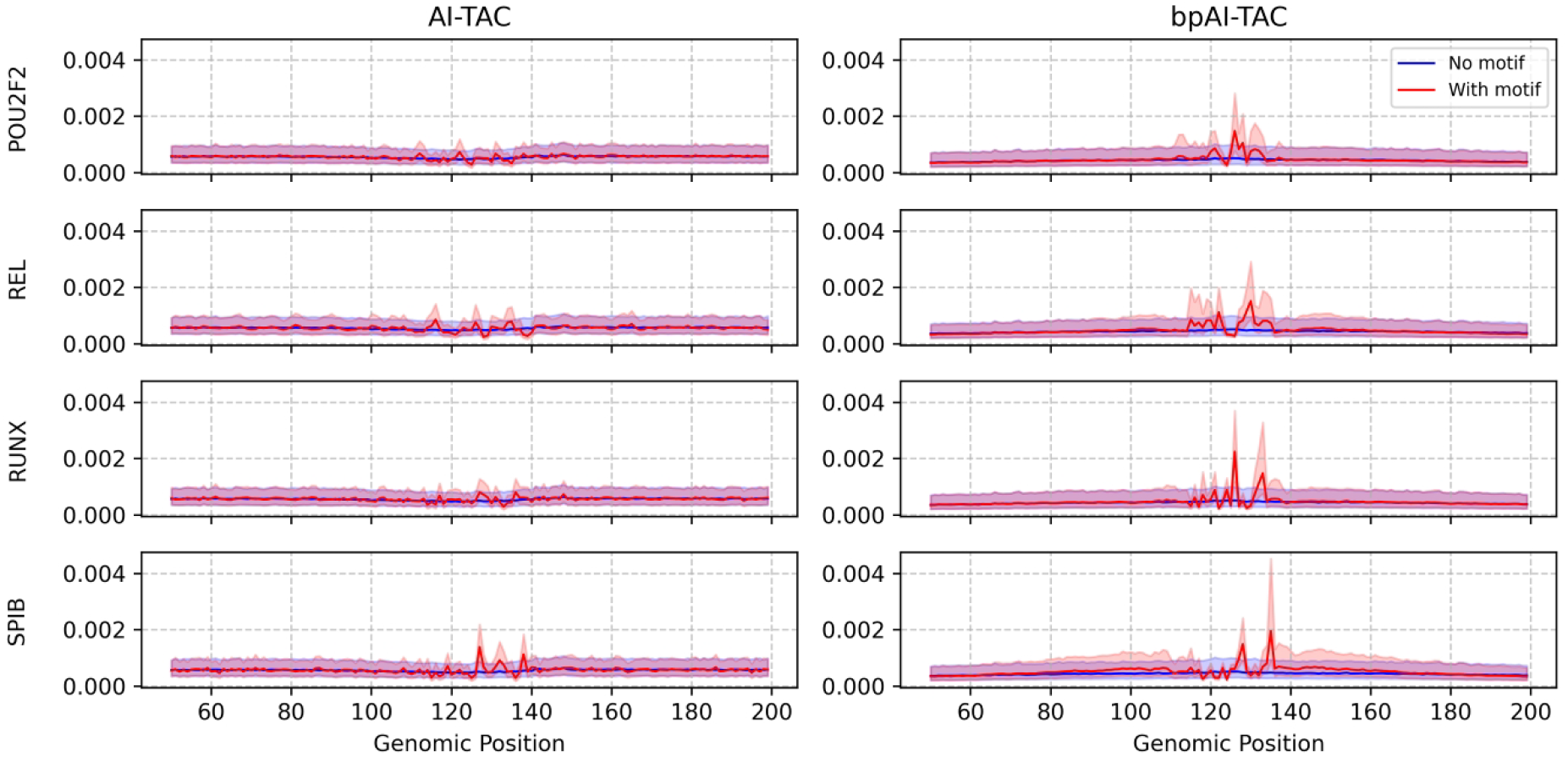
AI-TAC and bpAI-TAC Tn5 bias subtracted profile predictions on 4000 dinucleotide shuffled sequences with motifs inserted (red), and controls without any motifs inserted (blue). The red and blue lines correspond to the median predicted values, and the translucent bands correspond to the interquartile range. Predictions are averaged across five model initializations.

**Fig. S6.**
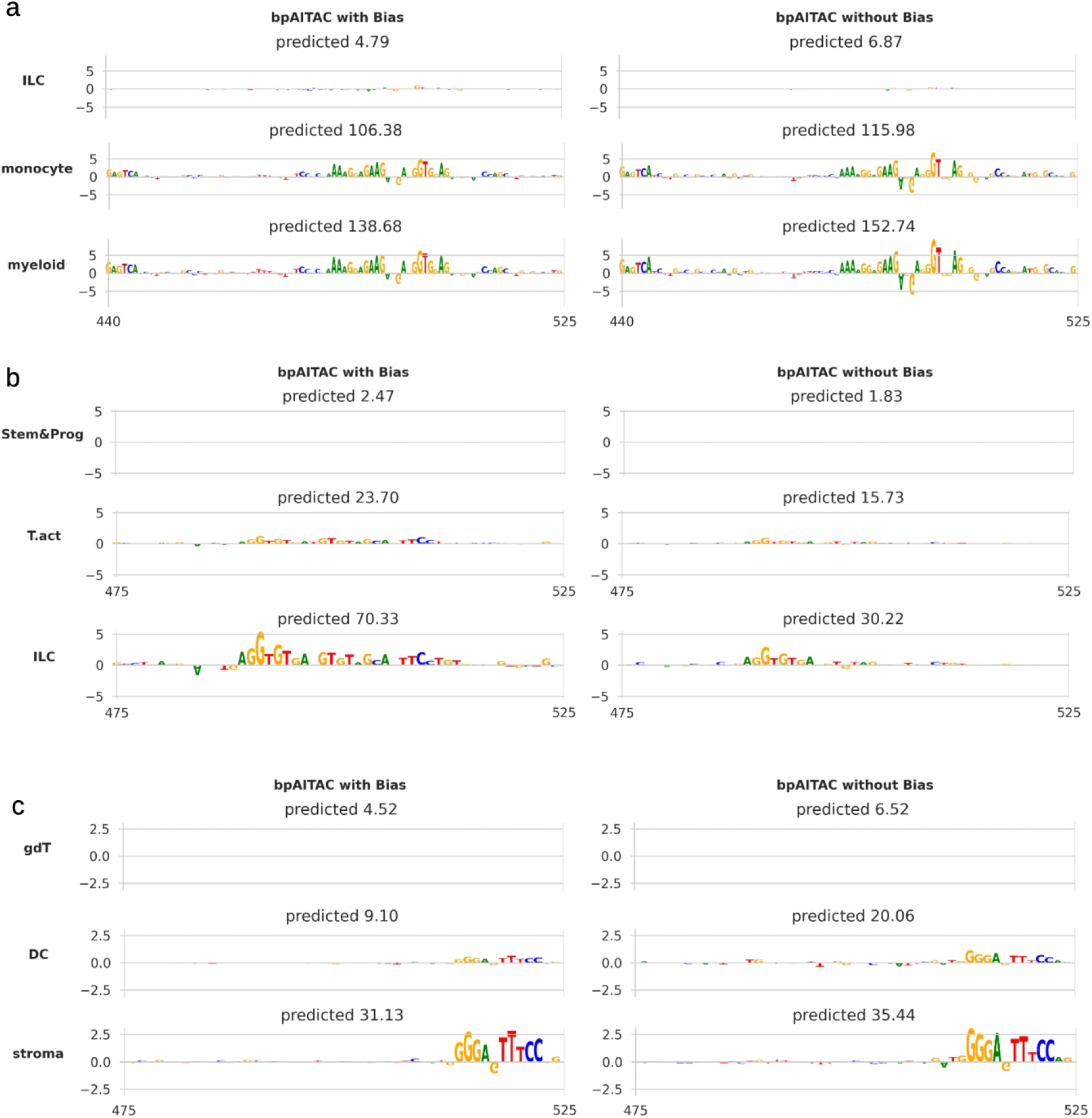
Tn5 bias correction shows no major effect on bpAI-TAC’s attribution maps for aggregated chromatin accessibility. Examples of bpAI-TACs attributions for chromatin accessibility of three regions, trained with and without the bias model. Attributions for the accessibility only show what contributes to the total number of Tn5 insertions, not the sequence elements that contribute to the shape of the profile.

